# Structural and biophysical properties of FopA, a major outer membrane protein of *Francisella tularensis*

**DOI:** 10.1101/2022.04.08.487631

**Authors:** Nirupa Nagaratnam, Jose M. Martin-Garcia, Jay-How Yang, Matthew R. Goode, Gihan Ketawala, Felicia M. Craciunescu, James D. Zook, Thomas D. Grant, Raimund Fromme, Debra T. Hansen, Petra Fromme

## Abstract

*Francisella tularensis* is an extremely infectious pathogen and a category A bioterrorism agent. It causes the highly contagious zoonosis, Tularemia. Currently, FDA approved vaccines against tularemia are unavailable. *F. tularensis* outer membrane protein A (FopA) is a well-studied virulence determinant and protective antigen against tularemia. It is a major outer membrane protein (Omp) of *F. tularensis*. However, FopA-based therapeutic intervention is hindered due to lack of complete structural information for membrane localized mature FopA. In our study, we established recombinant expression, monodisperse purification, crystallization and X-ray diffraction (∼6.5 Å) of membrane localized mature FopA. Further, we performed bioinformatics and biophysical experiments to unveil its structural organization in the outer membrane. FopA consists of 393 amino acids and has less than 40% sequence identity to known bacterial Omps. Using comprehensive sequence alignments and structure predictions together with existing partial structural information, we propose a two-domain organization for FopA. Circular dichroism spectroscopy and heat modifiability assay confirmed FopA has a β-barrel domain consistent with alphafold2’s prediction of an eight stranded β-barrel at the N-terminus. Small angle X-ray scattering (SAXS) and native-polyacrylamide gel electrophoresis revealed FopA purified in detergent micelles is predominantly dimeric. Molecular density derived from SAXS at 31 Å shows putative dimeric N-terminal β-barrels surrounded by detergent corona and connected to C-terminal domains via flexible linker. Disorder analysis predicts N- and C-terminal domains are interspersed by a long intrinsically disordered region and alphafold2 predicts this region to be largely unstructured. Taken together, we propose a dimeric, two-domain organization of FopA in the outer membrane: the N-terminal β-barrel is membrane embedded, provides dimerization interface and tethers to membrane extrinsic C-terminal domain via long flexible linker. Structure determination of membrane localized mature FopA is essential to understand its role in pathogenesis and develop anti-tularemia therapeutics. Our results pave the way towards it.

## Introduction

The life-threatening disease tularemia is caused by the extremely infectious intracellular pathogen, *Francisella tularensis* (1). It is a vector-born zoonosis (2) and endemic throughout the Northern Hemisphere (3). Insect bites, skin contact with contaminated materials, ingestion of infected animals, drinking contaminated water and inhaling contaminated air can cause human infections (4). According to the United States Centers for Disease Control and Prevention (CDC), approximately 100 human cases are reported every year in the United States. *F. tularensis* has low infectious doses (10-50 colony forming units) and can be easily disseminated by air (5). If untreated or misdiagnosed, tularemia can cause up to 30-60% fatality rates and therefore has significant potential to cause severe morbidity and mortality (6, 7). Due to these reasons, *F. tularensis* is designated as category A bioterrorism agent by the CDC.

So far, a United States Food and Drug Administration (FDA) approved vaccine against tularemia is not available. One of the most promising strategies to develop safe and effective vaccines employs acellular subunits, which are either recombinantly synthesized components or purified immunodominant antigens from the pathogen (8, 9). Over the past 15 years, research has focused on utilizing *F. tularensis* lipopolysaccharides (10-13) and outer membrane proteins (14-17) as subunit vaccine candidates. One such evidence-based, extensively studied subunit vaccine candidate is *F. tularensis* outer membrane protein A (FopA) (17). FopA is highly immunogenic (18-21) and is an abundantly expressed outer membrane protein of *F. tularensis* (22). Several studies have shown that FopA is a protective antigen against tularemia. Monoclonal antibodies against FopA partially protected mice infected with the *F. tularensis* live vaccine strain (LVS) (23). Administration of purified recombinant FopA protected mice challenged with lethal LVS (24). In addition, a cocktail of recombinant peptides made of epitopes from FopA and *F. tularensis* lipoprotein Tul4 elicited strong immunogenicity *in vivo* and *in vitro* (25, 26).

However, the precise biological function of FopA is not yet elucidated. In Gram-negative bacteria, the outer membrane and the proteins embedded into it directly interact with the external environment. Outer membrane proteins serve well-established roles in bacterial adhesion, invasion, and evasion of host immune responses (27). Emerging evidence shows that outer membrane proteins are necessary for the intracellular survival of many pathogens and act as virulence determinants for many diseases (28-30). Current knowledge of the molecular and genetic mechanisms of *F. tularensis* infectivity, evasion of host defense and virulence is still limited (31, 32). FopA is an important virulence determinant of *F. tularensis* as it is required for the intracellular proliferation of the fully virulent SCHU S4 strain in bone marrow derived macrophages and is essential for virulence in mice (33).

So far, three-dimensional structure of neither the full-length nor the transmembrane domain of FopA is available. Only the structure of membrane extrinsic domain of FopA has been recently determined by Michalska and co-workers and deposited in the Protein Data Bank (PDB ID 6U83) without a detailed publication. To advance the current knowledge of FopA immunogenicity and its role in *F. tularensis* virulence, structural information is essential, which is vital towards the rational design of effective drugs and subunit vaccines against tularemia. In the present study, as a pre-requisite for structure determination, we established high-level membrane directed recombinant expression and monodisperse purification of the membrane translocated mature FopA. Using a myriad of bioinformatics and biophysical techniques, crucial insights into the structure of FopA has been elucidated in detail and set the stage for the determination of first membrane-integral structure of FopA.

## Results

### FopA shares structural characteristics of OmpA family proteins

FopA is classified as an OmpA family protein according to the Pfam database (34) of protein families. To investigate OmpA family characteristics of FopA, a comprehensive bioinformatics analysis was carried out. Using NCBI’s protein BLAST server, outer membrane proteins (OMPs) that are homologous to FopA in sequence were identified. The closest homologs were putative OmpA family proteins that shared 26 – 40 % sequence identity with FopA (S1 Table). The seven most closely related proteins, based on the query cover, E (expect) value and percent identity, were chosen for multiple sequence alignment. These selected proteins were from organisms that all belong to the class Gammaproteobacteria. The alignment revealed two conserved regions localized to the N- and C-terminus. However, the homology is highest and most continuous in the C-terminal region (S1 Fig). The finding suggests that these two regions could fold into two individual domains.

Consistent with above finding, the NCBI’s conserved domain search predicted two putative conserved domains in FopA: a transmembrane β-barrel domain at the N-terminus and an OmpA-like domain at the C-terminus. The N-terminal domain (FopA-NTD) shares low sequence conservation with OmpA family proteins as described above (S1 Fig) and more importantly, an experimentally solved molecular structure of this domain is unavailable. Therefore, structure prediction was performed using aphafold2, the highly accurate deep learning based protein structure prediction algorithm (35). The predicted structure of FopA-NTD spans amino acids 75-210 (Fig 1A and 1B) and displays an anti-parallel eight stranded β-barrel with four large loops and three shorter turns (Fig 1B). On the other hand, the C-terminal domain of FopA (FopA-CTD) shares significant sequence conservation with OmpA family proteins as described above (S1 Fig) and has 3D-structure available in the protein data bank (PDB ID 6U83; unpublished) (Fig 1B). The structure of FopA-CTD spans amino acids 240-393 (Fig 1A and 1B). A structure-based sequence alignment of FopA-CTD with related OMPs of known structure was performed to identify regions with conserved secondary structural elements (S2 Fig). Comparison of FopA-CTD structure with other OmpA family proteins revealed that the secondary structure is conserved, where FopA-CTD adopts the characteristic βαβαββ topology of OmpA-like domains (36). Next, prediction of intrinsically disordered elements present within FopA was performed. The analysis revealed that the 29 amino acids long linker (aa 211-239; Fig 1A and 1B) between FopA-NTD and -CTD is highly disordered (Fig 1C), hence, very likely to be flexible. Additionally, this linker region exhibits no sequence conservation among the closest OmpA family homologs (S1 Fig). The tendency of this long segment to possess disorder was further investigated by analyzing the amino acid composition using ExPASy ProtParam (37), which showed this region lacks order promoting residues C, L, F, W and Y (38) (Fig 1D). The results are consistent with Alphafold2 which couldn’t predict defined structural elements within this region (Fig 1B).

**Fig 1.**
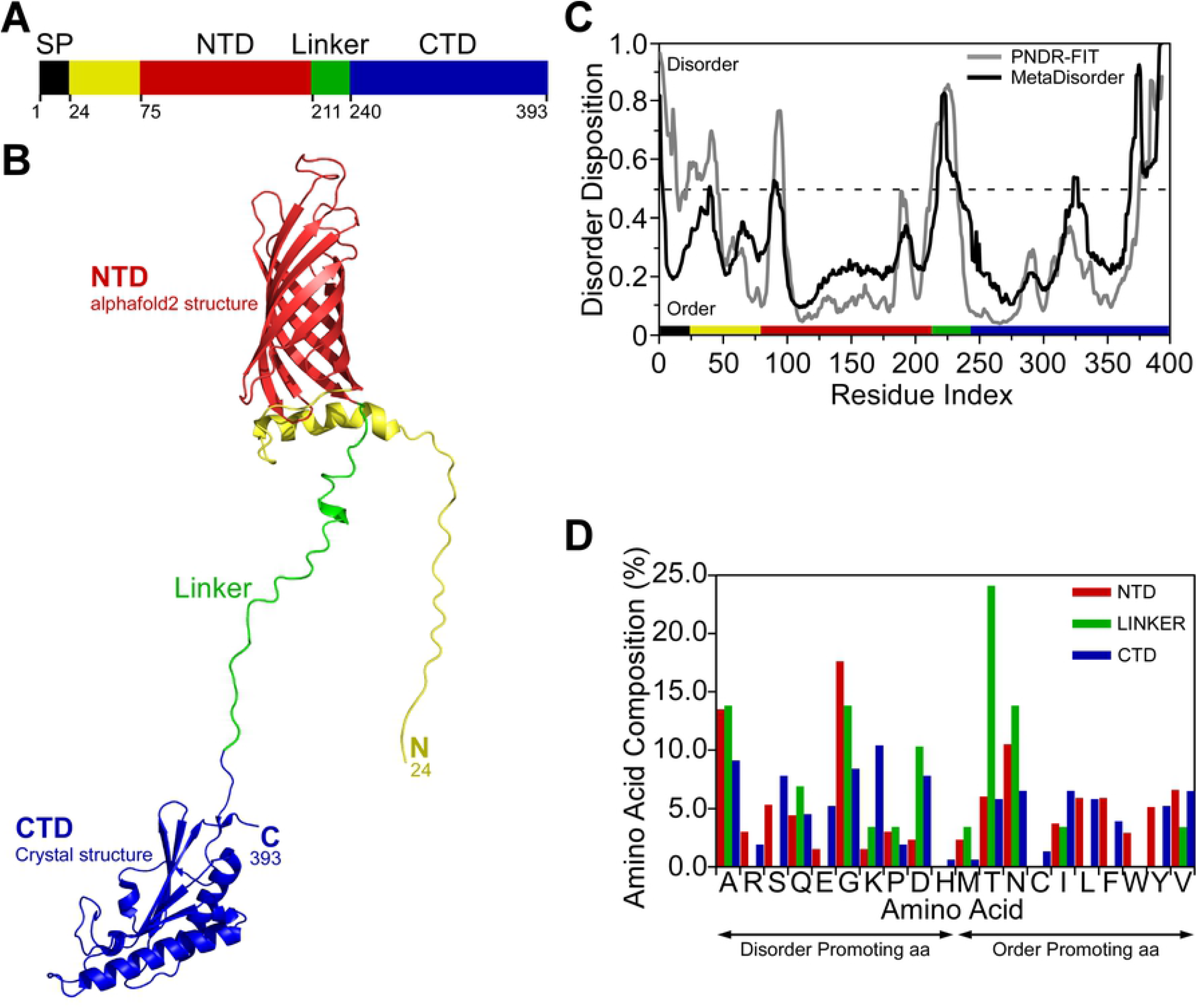
Structural analysis of FopA. (A) Domain organization of FopA. The colors of the domains are maintained in the subsequent panels. SP, signal peptide; NTD, N-terminal domain; CTD, C-terminal domain. (B) Alphafold2 model of FopA-NTD (red) and crystal structure of FopA-CTD (blue) solved by Michalska and co-workers (PDB ID 6U83). The unstructured linker connecting the two domains is shown in green. (C) Prediction of disordered regions in FopA by two meta disorder prediction programs PONDR-FIT (63) and MetaDisorder (60). (D) Comparison of the composition of each amino acid in the NTD, CTD and linker. Order and disorder promoting amino acids (aa) are indicated (38).

### Recombinant FopA is successfully expressed and translocated to *E. coli membranes*

Native FopA is a 41.4 kDa protein comprised of 393 amino acids. For crystal structure determination, producing high levels of FopA that is well folded during expression and purified to homogeneity is challenging. In our study, full-length FopA was recombinantly expressed in *E. coli* and the membrane translocated mature form was purified. Mature FopA referred to here and throughout the text corresponds to FopA with its signal peptide cleaved upon membrane translocation. To achieve successful membrane translocation, the native signal peptide of FopA (aa 1-23) (Fig 1A) was retained in the expression construct (Fig 2A). The *E. coli* protein translocation machinery recognizes this sequence and directs FopA to the cell membrane. SDS-PAGE of membrane fractions shows a putative FopA band in the outer membrane fraction (Fig 2B, lane OM) at the expected theoretical MW of mature FopA-His8 (40.2 kDa). Anti-His immunoblot confirms that FopA is located in the outer membrane (Fig 2C, lane OM), as well as in the inner membrane (Fig 2C, lane IM). In the subsequent experiments, this membrane translocated mature FopA is purified and characterized. Additional smaller bands are visible in some of the lanes (Fig 2C), which may be truncation products resulting from proteolytic degradation.

**Fig 2.**
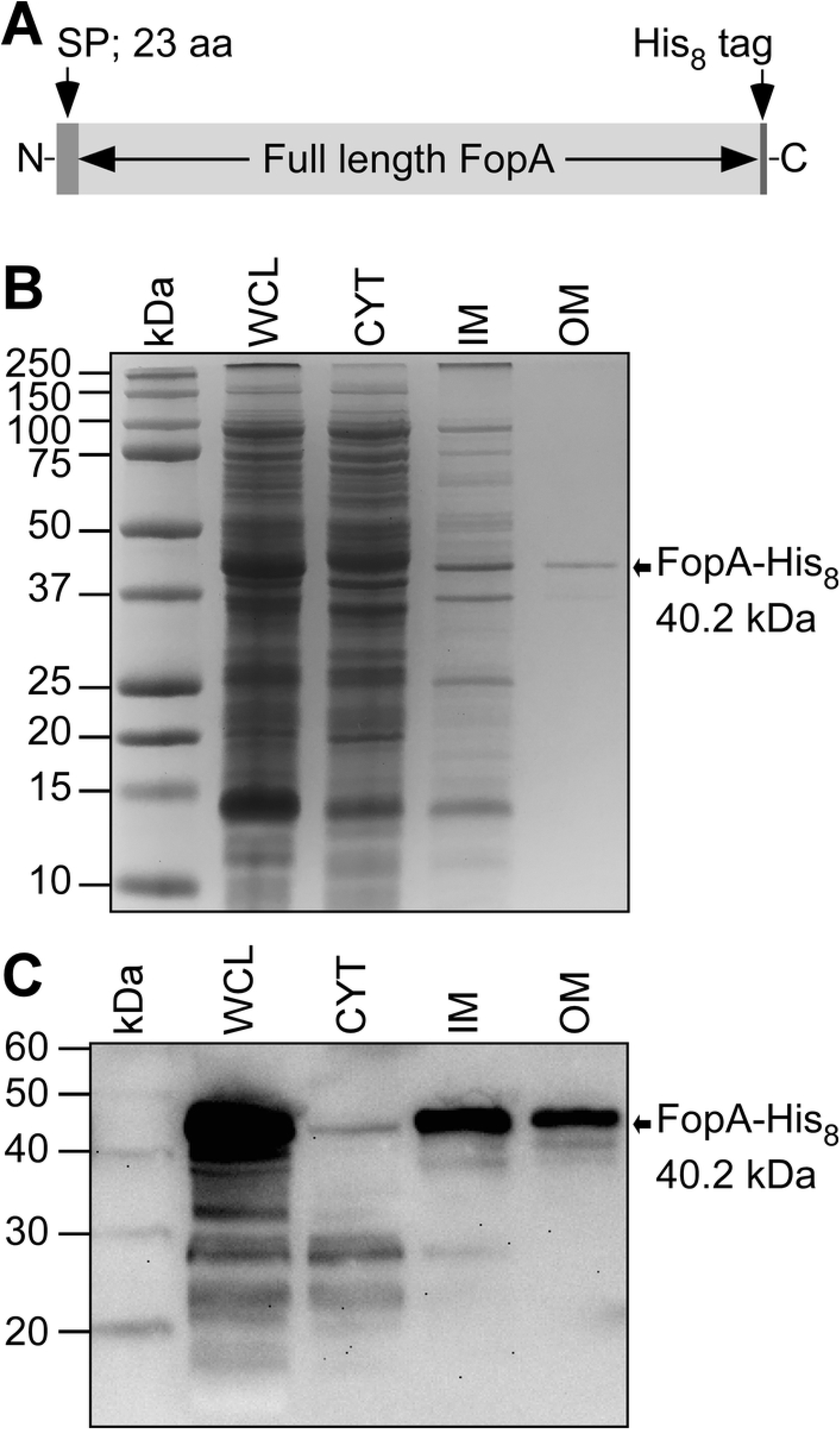
Subcellular localization of heterologously expressed FopA in *E. coli*. (A) Schematic of the FopA expression construct. Residues 1–23 are the signal peptide (SP). (B) SDS-PAGE of *E. coli* subcellular fractions. WCL, whole cell lysate; CYT, cytosol; IM, inner membrane; OM, outer membrane. The bands are visualized by Coomassie stain. (C) Immunoblot using anti-His antibody of the same samples as in B.

### Membrane localized mature FopA is detergent solubilized and extracted in large quantities

In order to efficiently detergent solubilize *E. coli* membranes and extract the membrane embedded FopA in the form of protein-detergent micelle, a detergent screen was performed for the isolated cell membranes using ten detergents: βDDM, βDM, OG, CHAPS, FC-12, LDAO, Cymal-2, Cymal-3, Cymal-4 and Cymal-6. Of these detergents, βDDM exhibited high degree of membrane solubilization, as revealed by anti-FopA immunoblot (S3 Fig, lane β-DDM), thereby extracting the vast majority of FopA into detergent micelles. Following optimizing the solubilization condition for βDDM, which includes systematic variation of detergent concentration, temperature and duration of solubilization, FopA was successfully purified from *E. coli* membranes. Briefly, the purification included βDDM extraction of the membrane fraction followed by Ni-NTA affinity chromatography and size exclusion chromatography (SEC) (Fig 3A and 3B). One of the major obstacles in obtaining highly pure FopA was the co-purification of FopA truncation products that result from proteolytic degradation. As predicted by bioinformatic analyses, FopA harbors long flexible regions (Fig 1B) making it more prone to proteolytic cleavage. The longer the detergent solubilization of *E. coli* membranes, the higher the proteolytic cleavage observed in purified FopA. This makes large scale purification of high-quality FopA for downstream biophysical and structural analyses challenging. SEC showed two main peaks, which were collected as pool B and C as shown in Fig 3B. Coomassie-stained SDS-PAGE (Fig 3C) showed that mature FopA was in pool B (lane B). The identity of this band as FopA was confirmed by anti-FopA immunoblot (Fig 3D, lane B). SEC successfully separated the lower molecular weight fragments into pool C (Fig 3C and 3D, lanes C).

**Fig 3.**
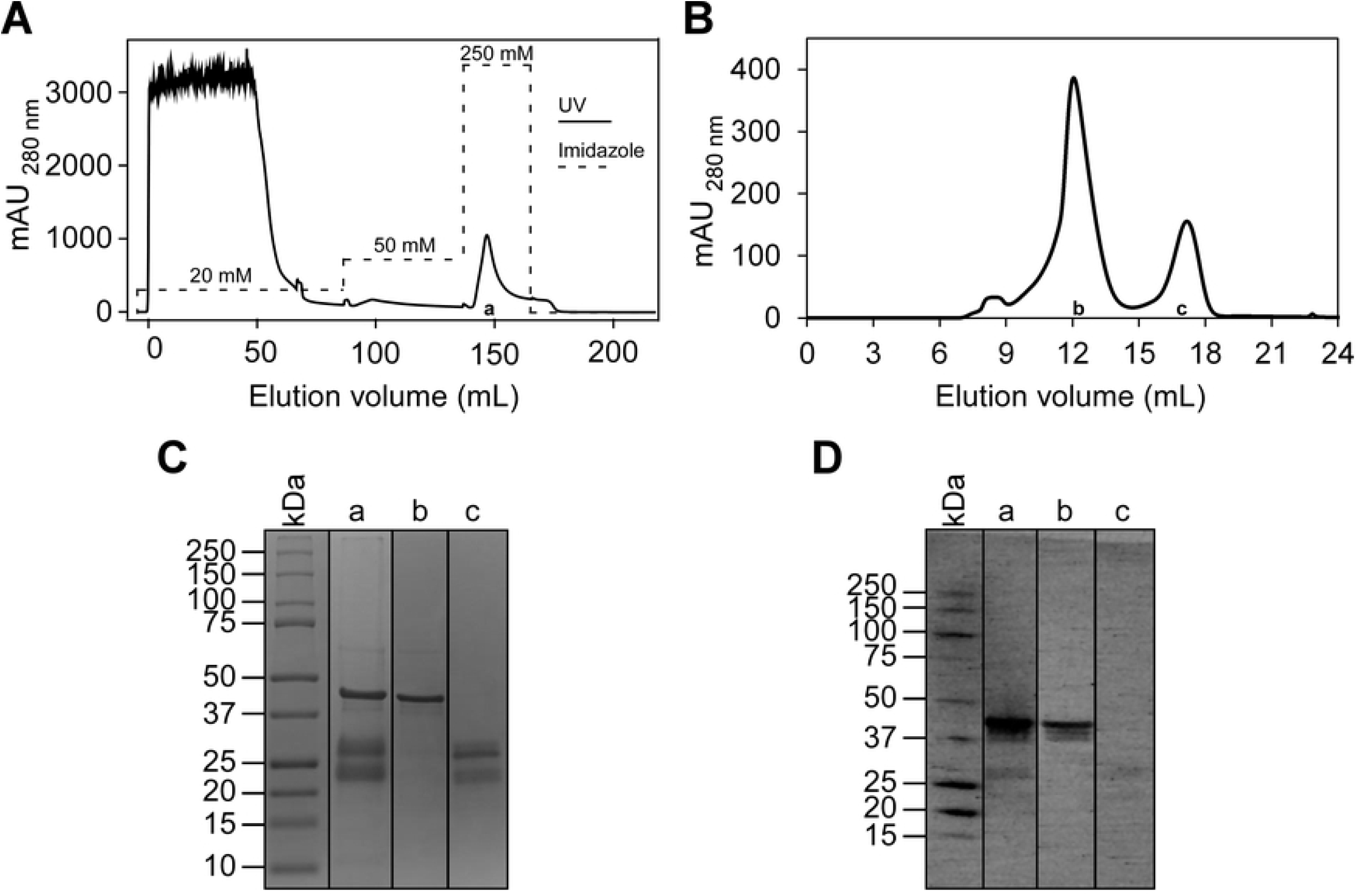
Purification of membrane translocated mature FopA. (A) The Ni-NTA elution profile and (B) SEC profile of FopA. (C) SDS-PAGE of FopA purification products visualized by Coomassie stain. Lane a, pooled elution fractions of FopA from Ni-NTA affinity chromatography; Lane b, pooled elution fractions of mature FopA (elution volume 10-14 mL) in SEC; lane c, pooled elution fractions of truncation products of FopA (elution volume 16-17 mL) in SEC. (D) Immunoblot using anti-FopA polyclonal antibody of the same samples as in C.

### FopA extracted into detergent micelles is further purified to highest homogeneity

During some of the purifications, the SEC peak that contained mature FopA at the elution volume (V_e2_) ∼12.5 mL, was preceded by a shorter shoulder peak at V_e1_ ∼10 mL (Fig 4A), indicating minor levels of heterogeneity in the purified FopA. This heterogeneity was further investigated by performing clear native PAGE on the peak fractions (V_e1_ ∼10 mL and V_e2_ ∼12.5 mL) of SEC (Fig 4B). While the shorter shoulder peak (V_e1_ ∼10 mL; Fig 4A) contained different multimers of FopA (Fig 4B, lane 1), the major homogeneous peak (V_e2_ ∼12.5 ml; Fig 4A) contained predominantly the dimeric form, based on its migration at an apparent MW of ∼80 kDa (Fig 4B, lane 2). In order to obtain highly monodisperse FopA, the elution fraction containing the dimeric FopA was collected at the apex of peak 2 (V_e2_ ∼12.5 ml) of the first SEC (Fig 4A) and subjected to a second SEC (Fig 4C). The relative MW (*M*_r_) of the eluted dimeric FopA (V_e_ ∼12.5 mL) from the second SEC (Fig 4C) was determined by comparing the elution volume parameter *K*_av_ (partition coefficient) of FopA with the *K*_av_ values obtained for known MW standards (Fig 4D). The analysis yielded the *M*_r_ to be ∼180 kDa (Fig 4D). Since, the dimeric FopA-His8 is only 80.4 kDa, the additional molecular weight could be attributed to the βDDM detergent micelle surrounding the membrane spanning region of FopA. Dynamic light scattering (DLS) of the FopA purified by a second SEC revealed a homogeneous size distribution of a single type of species with a particle hydrodynamic radius of ∼8 nm (S4A Fig). DLS further confirmed that this purified FopA at the concentration of 11 mg/mL is devoid of aggregates which makes the protein preparations highly suitable for crystallization experiments. Consistent with DLS, negative stain electron microscopy also showed a homogeneous particle distribution of FopA and a particle size of ∼10 nm (S4B Fig).

**Fig 4.**
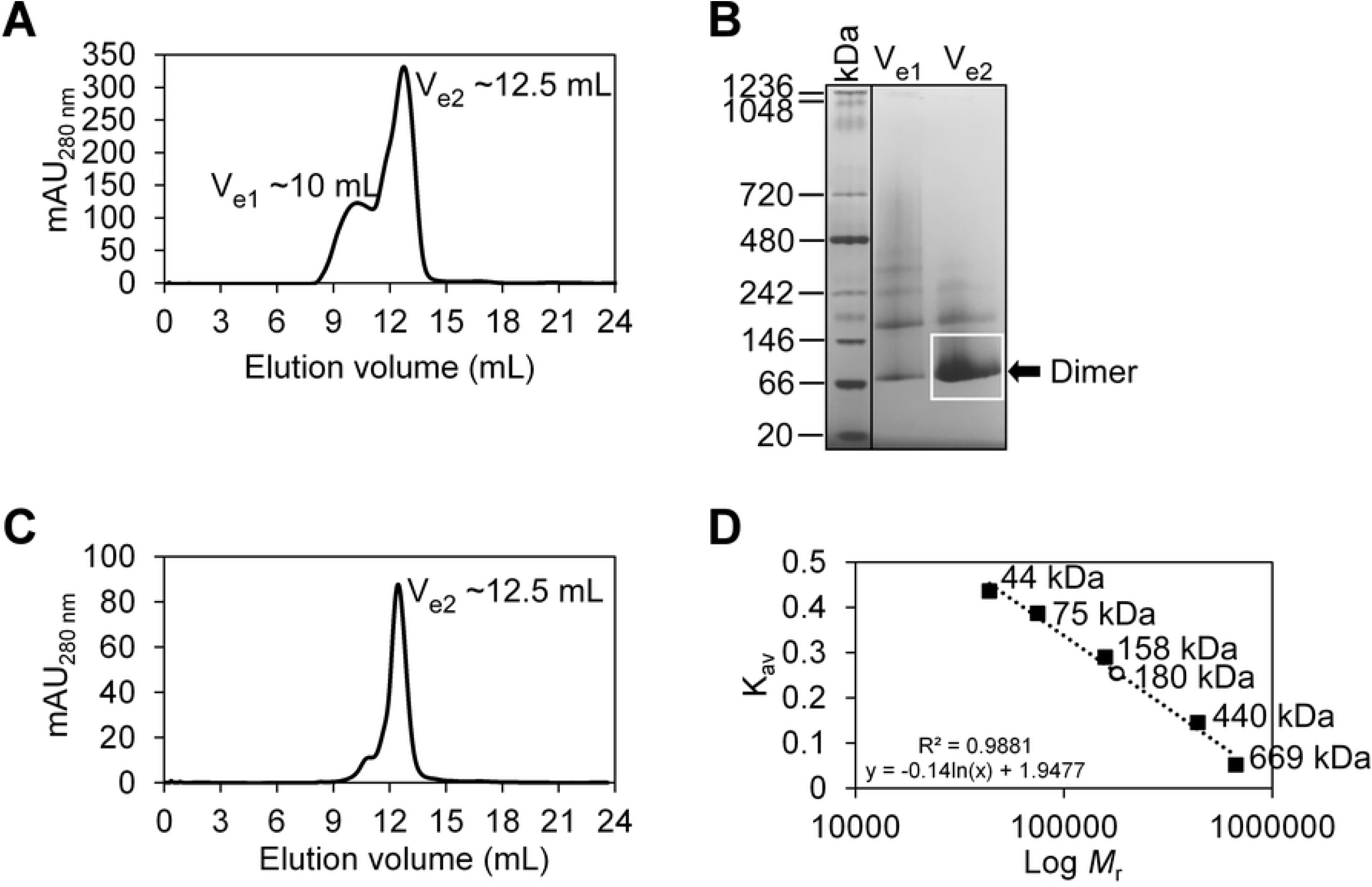
Assessing the oligomeric state and relative molecular weight (*M*_r_) of FopA. (A) Distribution of the various oligomeric states of FopA following first size exclusion chromatography (SEC). Elution peaks containing mature FopA are designated as V_e1_ and V_e2_. (B) Clear native-PAGE of the elution peaks of the first SEC. (C) Chromatogram of a second SEC that was loaded with peak V_e2_ from the first SEC in A. (D) SEC column calibration using known protein standards (black squares) and calculated *M*_r_ of dimeric FopA (open circle) eluted during the second SEC.

### FopA has a transmembrane β-barrel domain

Circular dichroism (CD) spectroscopy was performed to determine the secondary structural and folding properties of purified FopA. The far-UV CD spectrum indicated a pattern representative of β-strand-rich proteins, having minimum and maximum ellipticities near 218 and 196 nm, respectively (Fig 5A). The CD spectrum was further analyzed to estimate the secondary structural composition by the CDPro software package (39). The values obtained suggest FopA is comprised of 30.6% β-strands and 21.6% β-turns and were in accordance with secondary structure predictions made by bioinformatics analyses (S2 Table). Next, the β-barrel fold of FopA was assessed by heat modifiability assay coupled with SDS-PAGE. In the assay FopA was treated with SDS-PAGE Sample Buffer and incubated at different temperatures of 25°C, 37°C, 50°C, 75°C and 95°C. FopA which is completely unfolded by heat (95°C) migrated at the expected theoretical MW of mature FopA-His8 (40.2 kDa), while the non-heated (25°C), folded FopA migrated at an apparent MW of ∼35 kDa (Fig 5B). This characteristic migration difference indicates FopA is properly folded during recombinant expression. Additionally, the results show that the β-barrel fold is significantly heat-resistant and only begins to unfold above 75°C (Fig 4B).

**Fig 5.**
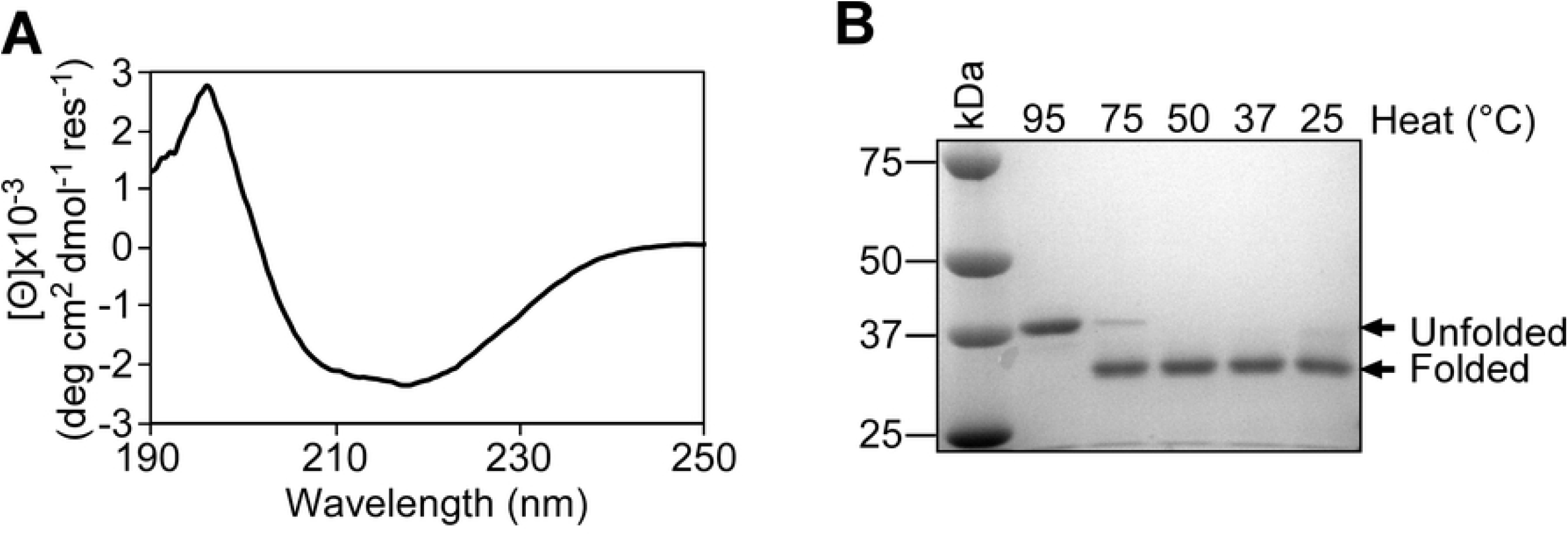
Secondary structural analyses of purified FopA. (A) Circular dichroism spectroscopy of membrane translocated FopA. The far-UV spectrum shows characteristic minimum and maximum ellipticities near 218 and 196 nm, respectively the characteristic values for β-strand-rich proteins. (B) Heat modifiability of FopA tested by SDS-PAGE stained with Coomassie blue. Unfolded FopA runs at the expected theoretical MW of mature FopA-His8 (40.2 kDa), while the folded FopA runs at an apparent MW of ∼35 kDa.

### Purified FopA is predominantly a dimer and has a distinct molecular envelope

Next, the highly homogeneous FopA was subjected to small angle X-ray scattering (SAXS). SAXS can be used to determine the molecular weight (MW), overall shape, and structural flexibility of a protein in solution. Guinier analysis of the FopA scattering data revealed a linear trend at low *q* values in the Guinier region (*q* < 1.3/*R*_*g*_; *R*_*g*_, radius of gyration) (Fig 6A and 6B), indicating that the analyzed FopA sample was devoid of aggregates. Radius of gyration (*R*_*g*_) estimated from the Guinier plot was found to be 44.0 +/-0.7 Å. The maximum dimension (*D*_*max*_) and the radius of gyration (*R*_*g*_) of FopA derived from the pair distance distribution function *P*(*r*) (Fig 6C) were ∼160 Å and 44.7 Å, respectively, in good agreement with the *R*_*g*_ found from the Guinier analysis. The left-skewed shape of the *P*(*r*) distribution (Fig 6C) indicates FopA is an elongated molecule. The Kratky plot shows a mostly folded protein with multiple peaks suggesting multiple domains connected by flexible linkers (Fig 6D). However, Kratky plots are difficult to interpret for membrane proteins with detergent. The presence of the detergent corona around FopA-NTD complicates the standard Kratky analysis, making it difficult to ascertain the state of flexibility directly from the plot.

**Fig 6.**
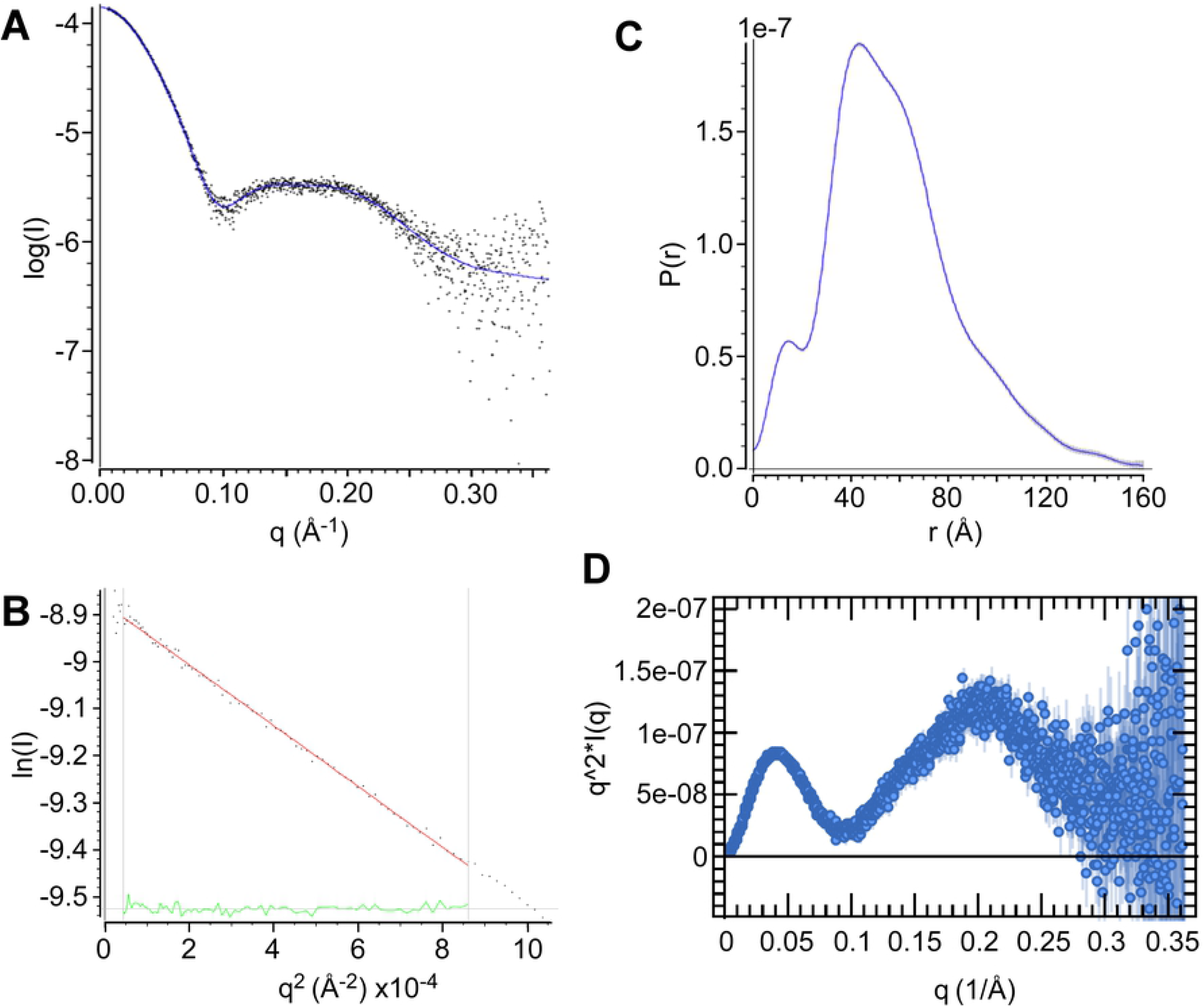
Small angle X-ray scattering of mature FopA. (A) Experimental scattering curve of FopA (black dots) and fit (blue line). (B) Guinier analysis in Primus (40). (C) Pair distance distribution function of FopA calculated by GNOM (41) from the experimental scattering curve. (D) Kratky plot with a characteristic peak suggesting correct folding, followed by a second peak which could imply either flexibility or effect of detergent belt surrounding the protein, a common signal in membrane protein analyses.

MW of FopA was estimated by three programs: GNOM in ATSAS (41), BioXTAS RAW (42) and DENSS (43) and using two methods Porod volume (MW_Porod_) (44) and Volume of correlation (MW_Vcorr_) (45) (Table 1). Five out of six alternate calculations yielded similar MW results of ∼84 kDa average. Overall, the results suggest FopA is present as a dimer in solution, which agrees with the theoretical MW of 80.4 kDa of dimeric mature FopA-His8.

**Table 1.** FopA MW estimation from SAXS data.

| Program | MW<sub>Porod</sub> (kDa) | MW<sub>Vcorr</sub> (kDa) |
| --- | --- | --- |
| GNOM in ATSAS | 193 | 88 |
| BioXTAS RAW | 74 | 89.3 |
| DENSS | 81 | 88 |

Electron density map at 31 Å (Fig 7) reconstructed by DENSS displays an overall cylindrical structure of FopA approximately ∼160 Å in the longest dimension and ∼90 Å along the remaining two axes. This observation is in good agreement with the *P*(*r*) distribution, which also indicated an elongated molecule with a maximum dimension of 160 Å (Fig. 6C). The FopA reconstruction exhibits high electron density in the center and at either end of the molecular envelope (Fig 7). The high density central core likely corresponds to the dimer of the N-terminal β-barrels, with the high density regions at either end possibly corresponding to the extended C-terminal domains connected by the flexible linker. The reconstruction also exhibits some small negative density regions immediately surrounding the high density core, possibly corresponding to the low-density detergent surrounding the transmembrane β-barrel domain.

**Fig 7.**
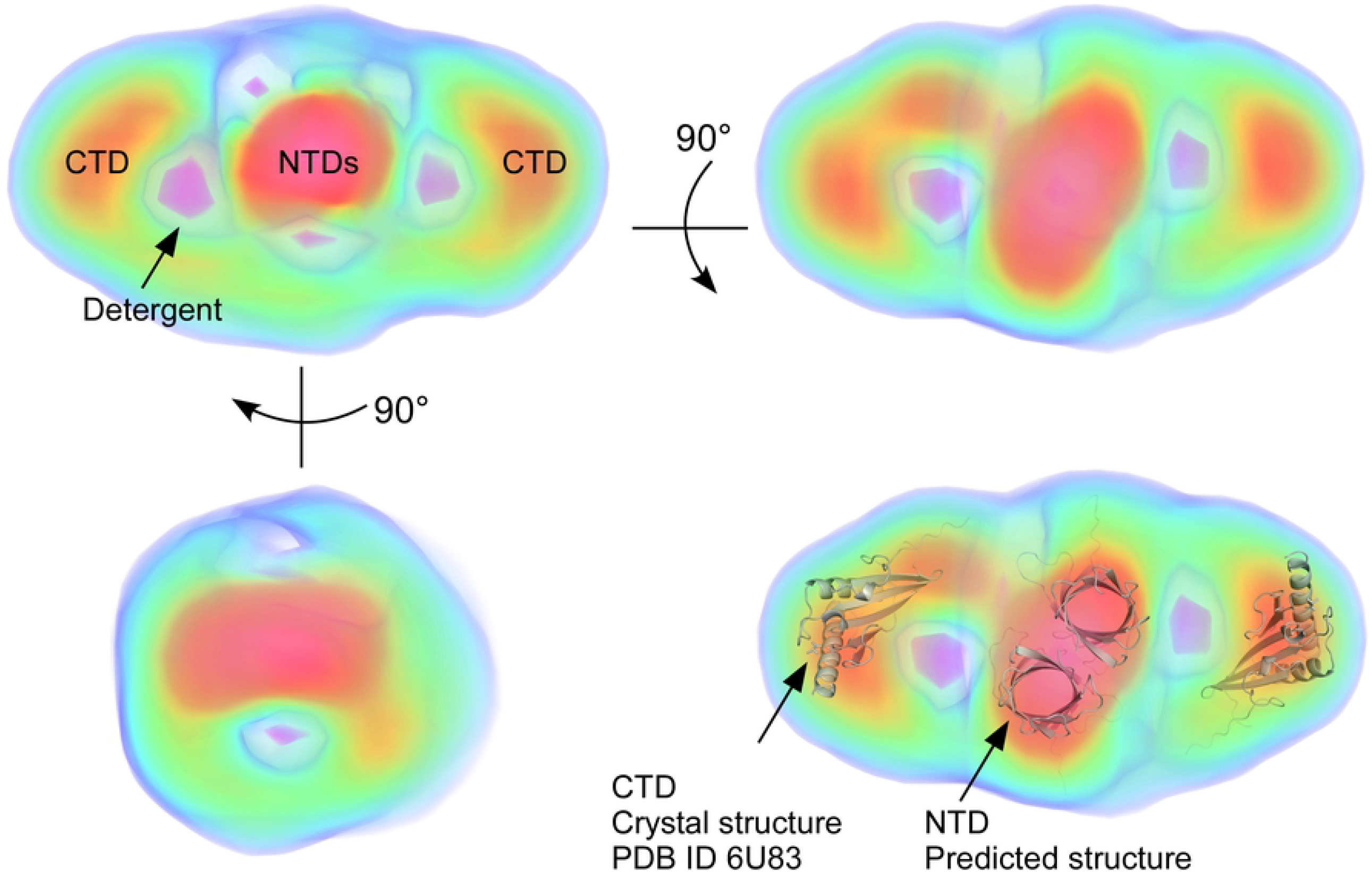
FopA electron density reconstructed by DENSS (43). Maps are colored according to density (blue=5 sigma, green=10 sigma, yellow=15 sigma, red=20 sigma, magenta=-5 sigma). The high density central core likely corresponds to the dimer of the N-terminal β-barrel domains (NTDs). The negative density (pink) surrounding the dimers belongs to low-density detergent corona. The high density regions at either end possibly corresponding to the extended C-terminal domains (CTD) connected by the flexible linker.

### FopA was crystallized and X-ray diffracted to ∼6.5 Å

Currently there is no structural information available for either the mature or the N-terminal transmembrane domain of FopA. Therefore, to obtain the first molecular structure of membrane translocated mature FopA, we pursued X-ray crystallography. The initial crystallization condition of FopA (27% 2-methyl-2, 4-pentanediol, 0.1 M Bis-Tris, pH 6.0, 1 mM CaCl_2_) was identified by crystallization screens using commercially available protein crystallization kits. Crystals were grown by hanging drop vapor diffusion with a protein to precipitant ratio of 1:1 (for further details see the Materials and Methods). Planar hexagonal crystals of size 25 × 25 × 1 μm (Fig 8A) were grown within 3 – 7 days. To verify these crystals are made of mature FopA, they were isolated, washed and analyzed by SDS-PAGE. The results confirmed that the crystals consisted of the mature FopA protein (Fig 8B). These crystals were directly frozen in liquid nitrogen and X-ray diffracted under cryogenic conditions at APS, Argonne National Laboratory. The crystals showed diffraction extending to ∼6.5 Å (Fig 8C). Even though these are very promising results as they show that mature FopA can be crystallized, the diffraction images revealed that the crystals feature a significant level of anisotropy. In addition, the diffraction resolution of the crystals had to be further improved.

**Figure 8.**
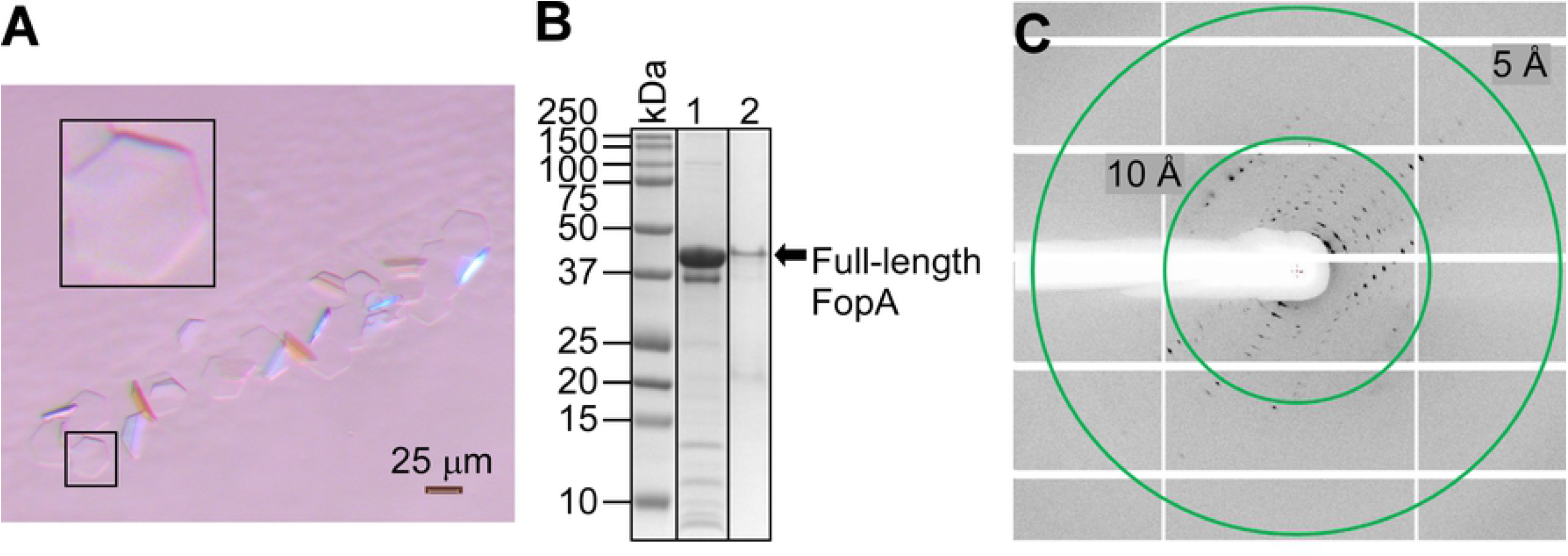
Crystallization of mature FopA and X-ray diffraction. (A) polarized light micrograph of FopA crystals. The larger black inset panel is an enlarged view of a planar hexagonal crystal that is shown in the small black inset panel. (B) SDS-PAGE of washed crystals. The mature FopA protein is present in the crystals as confirmed by the Coomassie-stained SDS-PAGE gel. Lane 1, mature FopA protein that was used in the crystallization; lane 2, mature FopA from isolated and washed crystals. (C) X-ray diffraction pattern from a single FopA crystal. Resolution rings are shown as green circles.

As the next step towards structure determination, the initial crystallization condition in Figure 8A was successfully reproduced and X-ray diffraction data collection was attempted. However, the planar dimensions and fragility of these crystals made harvesting and cryo cooling prior to X-ray diffraction extremely challenging. Therefore, high throughput crystallization screenings were performed to improve the shape, size, and diffraction quality of these crystals by varying salt and precipitant concentrations, incorporation of additives and changing pH values of the reservoir solution. From this fine screening of conditions, a total of 43 crystals were harvested from the reservoir solution, transferred to cryo protectant which had 25-30% glycerol and X-ray diffraction was performed under cryogenic conditions. Two of these crystals diffracted to a resolution of ∼6.5 Å and complete data set were collected. Unfortunately, the diffraction images still showed a similar pattern as described above with severe anisotropy and contained low number of unique reflections, which made structure determination of the FopA from the diffraction data extremely challenging. Nevertheless, the diffraction data were successfully indexed in H32 space group with unit cell parameters of *a* = 48.81 Å, *b* = 48.81 Å, *c* = 323.81 Å and α = 90°, β = 90°, γ = 120° (S3 Table). As discussed in more detail below we hypothesize that the highly flexible, long linker that connects the NTD and CTD of FopA may be attributable to the difficulties in obtaining high quality crystals for FopA.

## Discussion

This work reveals structural and biophysical properties of membrane translocated mature FopA. Although, FopA is classified as an OmpA family protein, FopA exhibits minimal overall sequence conservation (<40%) with OmpA family proteins from Gammaproteobacteria. OmpA family proteins are surface exposed porins present in the bacterial outer membrane in high-copy number and characterized by a two-domain structure: an N-terminal transmembrane β-barrel domain and a C-terminal periplasmic globular domain (27). Multiple sequence alignment of FopA with other OmpA family proteins shows two conserved clusters localized to the N- and C-terminal regions of FopA. These results suggest that the N- and C-terminal regions could independently fold into two individual structural domains. The intervening stretch of ∼29 amino acids shows no sequence homology and is highly disordered as revealed by our disorder prediction analysis. In fact, NCBI’s conserved domain search further supports this notion by predicting the N-terminal region of FopA to be folded into a β-barrel domain while the C-terminal region could adopt an OmpA-like domain. OmpA-like domains are found in the C-terminus of many Gram-negative bacterial outer membrane proteins. These domains interact non-covalently with the peptidoglycan layer and display a characteristic βαβαββ architecture similar to the C-terminal domain of *E. coli* outer membrane protein OmpA (OmpA-CTD) (27, 36). The recently deposited crystal structure of the C-terminal domain of FopA (FopA-CTD) by Michalska and co-workers in the Protein Data Bank (PDB ID 6U83) is in good agreement with the reported structures of other OmpA-like domains of many outer membrane proteins as revealed by our structure-based sequence alignment. However, the structure of neither the membrane spanning N-terminal domain (FopA-NTD) nor the mature FopA has yet been experimentally determined. Therefore, detailed biophysical and structural investigations of membrane translocated mature FopA are crucial towards deciphering its role in *F. tularensis* virulence and towards the search for therapeutics against tularemia.

In order to obtain structural insights of FopA-NTD, structure prediction was performed using alphafold2 (35). The predicted structure of FopA-NTD displays a β-barrel fold which is consistent with our CD spectroscopy data. In addition, folding properties of purified OMPs can be evaluated by heat modifiability assay (46). Heat modifiability is referred to the difference in the migration of heated versus non-heated samples of β-barrel outer membrane proteins from Gram-negative bacteria in SDS-PAGE. The numerous hydrogen bonds which hold the β-strands together in the barrel structure resist denaturation by SDS alone at room temperature and maintains the folded state of OMPs. Thus, non-heated OMPs can adopt a more compact structure than the heated OMPs that are completely unfolded. Nano demonstrated that FopA is located in the outer membrane of *F. tularensis* and heat modifiable (20). Membrane translocated mature FopA purified and characterized in our studies shows excellent heat modifiability confirming FopA-NTD is correctly folded into a β-barrel structure during recombinant expression. Native-PAGE and SAXS experimental data show purified membrane translocated mature FopA predominantly exists as a dimer. Finally, the electron density seen in our SAXS molecular envelope supports a dimeric, two-domain conformation: the N-terminal β-barrels forms a dimer surrounded by detergent corona and the globular C-terminal domains are extended and connected to the β-barrels via unstructured flexible linker. In agreement with this, the Kratky plot of FopA obtained from the SAXS experiments shows characteristics of a multidomain protein with flexible regions. Additionally, in the SAXS envelope, the nature of the high density central core corresponding to the dimeric N-terminal β-barrels can hypothetically be explained as follows: it could either correspond to two individual β-barrels or one large single β-barrel formed from the two monomeric β-barrels.

Taken together, with the experimental evidence we obtained in this study for FopA along with the current knowledge of the structures of OmpA family proteins, we propose a tertiary conformation for membrane translocated mature FopA: the β-barrels of two NTDs provide dimerization interface and localize in the outer membrane, while the two FopA-CTDs reside in the periplasm. Further, the FopA-NTD and -CTD are connected by the long, flexible linker (Fig 9). Both the length and flexibility of the interdomain linker (>29 amino acids) should enable FopA-CTD to reach the peptidoglycan layer and provide flexible mechanical support to the cell wall, a similar role demonstrated for *E. coli* OmpA (47). Given that FopA is one of the most abundant outer membrane proteins in *F. tularensis*, its non-covalent interaction with the peptidoglycan layer should have a significant role in maintaining membrane integrity. In addition, in the dimeric model, we hypothesize that the two NTDs either can form two individual narrow pores (Fig 9, Model 1) or both NTDs may rearrange and refold into a single large pore (Fig 9, Model 2) and possibly have a role in solute transport across the outer membrane.

**Figure 9.**
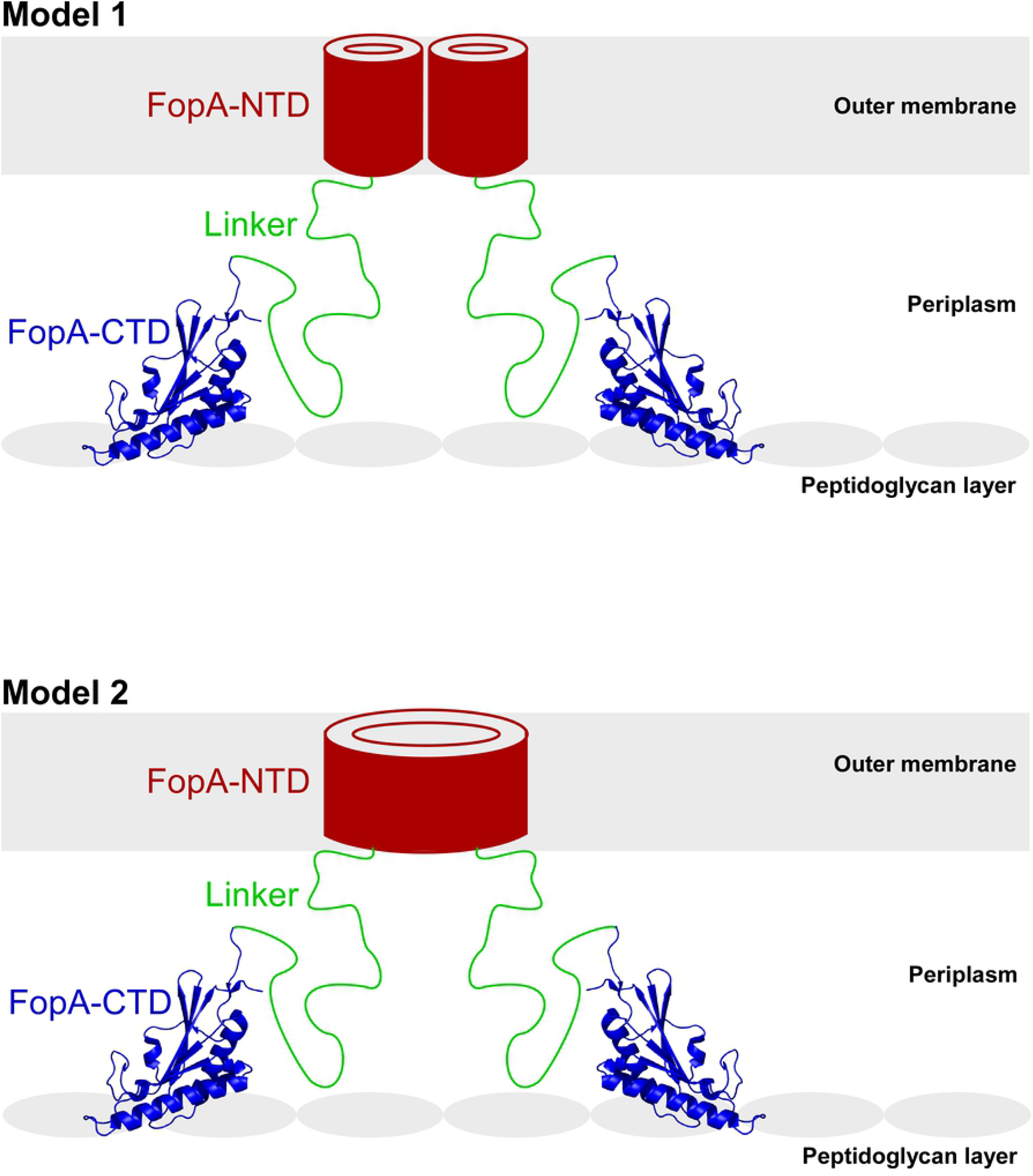
Proposed models of the structural organization of FopA in the outer membrane. FopA forms a dimer. FopA-NTD (red) folds into a β-barrel and is embedded in the outer membrane, FopA-CTD (blue; PDB ID 6U83) reside in the periplasm and interacts with the peptidoglycan layer and the long flexible linker (green) tethers both domains. **Model 1**: the two NTDs may form two individual narrow pores; **Model 2:** β-strands of both NTDs may rearrange and refold into a single large pore.

Finally, our goal to determine the high resolution structure of FopA by X-ray crystallography has not been reached yet, however we made progress towards it. For the first time, we demonstrated the crystallization of membrane translocated mature FopA. However, the long flexible linker, which is predicted to be largely disordered, is highly likely to adversely affect the diffraction quality of the current FopA crystals. Albeit this, the crystals diffracted to 6.5 Å resolution enabling the collection of the first complete X-ray data set. The diffraction data set was successfully indexed, and preliminary unit cell parameters were obtained. However, low resolution, anisotropy and low number of unique reflections in our X-ray diffraction data set posed challenges in solving the molecular structure of FopA. Improving the diffraction quality of FopA crystals is in progress and may require modifications to the flexible linker. Overall, with the procedures now established for expression, purification, biophysical characterization, and crystallization, we have set a clear path towards the structure determination of membrane translocated mature FopA. Determining the structure of FopA from *F. tularensis* should provide crucial insights into its virulence mechanism, pore formation and solute transport and to the overall architecture of this fascinating outer membrane protein.

## Materials and Methods

### Bioinformatics analyses of FopA

Pfam 32.0 database of protein families (48) was used to analyze FopA domain organization. Presence of a signal peptide in FopA was predicted by SignalP-5.0 (49). Proteins homologous to FopA sequence were identified by BLAST, and multiple sequence alignment was performed using the program M-Coffee (50). Sequence similarities of the aligned sequences were analyzed by the similarity depiction program ESPript 3.0 (51). Structure-based sequence alignment with homologous sequences of known structures was performed by ENDscript 2.0 (51). FopA structure prediction was performed by alphafold2 (35). FopA disorder prediction was performed by two meta-predictors PONDR-FIT (52) and MetaDisorder (53). Amino acid composition of FopA was calculated by ExPASy-ProtParam (37).

### Plasmid construction and transformation of *E. coli*

All chemicals used in this study were purchased from Millipore Sigma, unless otherwise noted. The *fopA* gene (FTT0583) from *F. tularensis* subsp. *tularensis* strain SCHU S4 was cloned into the *Bse*RI sites of the vector pRSET-C8xHis (DNASU Plasmid Repository accession number EvNO00629010) to generate the expression construct pRSET-FTT0583-His8 (DNASU FtCD00697191), which contains an octa-Histidine tag at the C-terminus. The membrane targeting native signal peptide sequence of FopA was retained in the coding sequence of the *fopA* gene during cloning (S5 Fig). The expression construct was transformed (54) into the *E. coli* C43(DE3) expression strain and selected for transformants on LB (Luria-Bertani) agar plates with 50 µg/mL carbenicillin (Teknova).

### FopA expression in *E. coli*

All bacterial growths were performed in a rotatory shaking incubator (InnovaTM 4230, New Brunswick Scientific) at 250 RPM. A single colony from the freshly transformed LB plate was inoculated into 10 mL of TB (Terrific broth) medium and grown overnight at 37 °C to obtain a starter culture. 1000 mL of TB supplemented with 50 μg/mL carbenicillin was inoculated with 10 mL of the overnight starter culture and grown at 37°C until the OD_600_ reached ∼0.5-0.7. Next, protein expression was induced with 1 mM IPTG (isopropyl β-D-1-thiogalactopyranoside; GoldBio) followed by expression for 17 hr at 25°C. Cells were harvested by centrifugation at 7,500 x g for 20 min at 4°C and frozen as a dense cell pellet at −80°C until use.

### *E. coli* subcellular fractionation

All centrifugations were done at 4°C. The frozen cell pellet from 1 L culture was re-suspended in PBS (phosphate buffered saline; 10 mM Na_2_HPO_4_, 1.76 mM KH_2_PO_4_, 136.89 mM NaCl, 2.68 mM KCl), pH 7.4, supplemented with 30 U/mL benzonase and EDTA-free protease inhibitor cocktail tablet (one tablet per 10 g of cell pellet). Cells were lysed by sonication (Branson 550 Sonifier with 1/2” Horn) using 3 cycles with 1 min cooling period on ice between cycles. Each cycle consisted of sixty 1 s pulses at 50% amplitude. The cell lysate was centrifuged at 10,000 x g for 20 min to separate unbroken cells and cell debris. The supernatant containing the cytosol and membrane fragments was centrifuged at 100,000 x g for 1 h. The pellet containing both inner and outer membrane fragments was re-suspended in PBS, pH 7.4, supplemented with protease inhibitors. To solubilize inner membrane fragments, 0.5% *N*-lauroyl sarcosine sodium salt was added to the membrane re-suspension and incubated for 2 hr with shaking at 4°C. The re-suspension was centrifuged at 100,000 x g for 1 h to pellet the outer membranes.

### FopA purification from *E. coli* membranes

All chromatography purification steps were carried out at 4°C using an ÄKTA pure fast protein liquid chromatography (FPLC) system (GE Healthcare). All buffer solutions were chilled prior to use. Cell lysis and membrane isolation were performed as described above with slight modifications. After separating unbroken cells and cell debris, the supernatant containing the cytosol and membrane fragments was centrifuged at 50,000 x g for 1 h. Next, the pelleted membrane fragments were re-suspended in PBS, pH 7.4 supplemented with protease inhibitors and 20 mM imidazole followed by detergent solubilization by adding 2% βDDM and incubating with shaking for 17 hr at 4°C. The detergent extract was centrifuged at 50,000 x g for 1 h. The supernatant containing detergent solubilized His-tagged FopA was filtered through a 0.22 μm cellulose acetate membrane filter (Foxx Life Sciences) and loaded at 1 mL/min onto a Ni-NTA column (HisTrap HP, 5 mL; GE Healthcare) that had been previously equilibrated with Ni-NTA Buffer (NB; 20 mM Bicine, pH 8, 500 mM NaCl, 20 mM imidazole, 0.05% βDDM) for affinity purification. After collecting the flow through, the column was washed with 20 column volumes of NB followed by 15 column volumes of NB supplemented with 50 mM imidazole. FopA was eluted with 5 column volumes of NB supplemented with 250 mM imidazole. Fractions containing FopA were concentrated up to 10-15 mg/mL using 100 kDa cut-off, 15 mL concentrators (Millipore) prior to further purification by size exclusion chromatography (SEC). SEC was performed using a Superdex 200 Increase 10/300 GL column (GE Healthcare) that was previously equilibrated in the SEC Buffer (20 mM Tris, pH 7.5, 150 mM NaCl, 0.05% βDDM). FopA was injected at 10-15 mg/mL in a 500 μL volume to the SEC column and eluted at 0.5 mL/min. Fractions containing FopA were stored at 4°C until further use. For some of the FopA purifications, a second SEC was performed under the same conditions as described above to obtain highly monodisperse FopA. Protein concentration was determined spectrophotometrically at 280 nm, using the extinction coefficient of 51,465 M^-1^ cm^-1^. The extinction coefficient and theoretical pI were calculated using ExPASy ProtParam (37).

### Detergent screening for *E. coli* membrane solubilization

Small-scale detergent screen was performed using ten different detergents: *n*-dodecyl-β-D-maltopyranoside (βDDM), *n*-decyl-β-D-maltopyranoside (βDM), *n*-octyl-β-D-glucopyranoside (OG) (GLYCON Biochemicals, GmbH), 3-[(3-cholamidopropyl)dimethylammonio]-1-propanesulfonate (CHAPS), Fos-Choline-12 (FC-12), lauryldimethylamine *N*-oxide (LDAO), 2-cyclohexyl-1-ethyl-β-D-aaltoside (Cymal-2), 3-cyclohexyl-1-propyl-β-D-maltoside (Cymal-3), 4-cyclohexyl-1-butyl-β-D-maltoside (Cymal-4), and 6-cyclohexyl-1-hexyl-β-D-maltoside (Cymal-6) (Anatrace). Briefly, *E. coli* cell membranes were isolated as described above and re-suspended in PBS, pH 7.4, supplemented with protease inhibitor. The membrane suspension was divided into ten portions of equal volumes and incubated with shaking with each of the ten detergents for 2 hr at 4°C, followed by centrifugation at 17,000 x g for 30 min at 4°C. Supernatants which contained the solubilized FopA as protein detergent micelles of corresponding detergent were analyzed by SDS-PAGE and immunoblotting.

### SDS-PAGE

SDS-PAGE was performed as reported previously (55) using Tricine SDS polyacrylamide gels with a 4 % stacking gel and resolving gels of 8 %, 10 % or 12 %. Protein samples were mixed with the SDS-PAGE Sample Buffer (10% glycerol, 60 mM Tris/HCl, pH 6.8, 2% SDS, 0.01% bromophenol blue and 1.25% beta-mercaptoethanol) in the volume ratio of 1:4 and heated at 95°C for 10 min prior to loading the gel. Protein bands were visualized by either staining with InstantBlue (Expedeon) Coomassie stain or were subjected to immunoblotting.

### Immunoblotting

Proteins were electro-transferred (Mini Trans-Blot Module, Bio-Rad) onto a PVDF membrane (Bio-Rad) for 1 h at 0.22 A using chilled Transfer Buffer (24.9 mM Tris base, 192 mM glycine, 20% (v/v) methanol). Hereafter, all steps were carried out at room temperature. Following transfer, membrane was blocked with PBS supplemented with 0.05% Tween 20 and 5% non-fat dry milk for 1 h with shaking. For anti-FopA immunoblotting, the membrane was subsequently incubated with polyclonal mouse anti-FopA antibody (Hansen *et al*., 2016) at 1:2000 dilution in the blocking solution for 1 h with shaking. The membrane was next washed with PBS supplemented with 0.05% Tween 20 for 20 min with shaking. The membrane was similarly incubated with Goat-anti-Mouse IgG (H+L) HRP conjugated secondary antibody (Invitrogen) at 1:3000 dilution and with the ladder conjugate (Precision Protein StrepTactin-HRP Conjugate, Bio-Rad) at 1:5000 dilution. Next, the membrane was washed as described above. To visualize the protein bands, chemiluminescent detection was performed using the Immun-Star HRP substrate kit (Bio-Rad) per manufacturer’s instructions. Anti-His Western blots were performed using Penta-His mouse monoclonal IgG^1^ as primary antibody (Qiagen, #34660) at 1:2000 dilution, HRP sheep-anti-mouse IgG as secondary antibody (Jackson ImmunoResearch, #515-035-062) at 1:5000 dilution and MagicMark XP Western Protein Standard (Life Technologies).

### Circular dichroism spectroscopy

Circular dichroism spectroscopy (CD) was performed on a JASCO J-710 CD Spectropolarimeter at 20°C. FopA was exchanged from SEC buffer to the CD Buffer (20 mM Tris, pH 7.5, 150 mM NaF, 0.05% βDDM) using 0.5 mL Amicon Ultra 100 kDa cutoff centrifugal filter unit (Millipore Sigma). Far-UV spectra (190–250 nm) were recorded using a bandwidth of 0.5 nm and response time of 4 s, in a 0.1 cm path-length quartz cuvette with 5 µM protein. Each spectrum was the average of five scans, with a scan rate of 50 nm/min. Spectra were corrected for background solvent effects by subtracting the signal from the buffer and then normalized as mean residue molar ellipticity. The molar ellipticity in deg. cm^2^/dmol was calculated as described (56). The spectrum was deconvoluted to estimate the protein secondary structure content using CDPro (39).

### Clear native-PAGE

Clear native-PAGE was performed using 4 - 16 % Bis-tris gels (Novex, Life Technologies), and NativeMARK unstained ladder was used as size standards (Thermo Fisher Scientific). Prior to loading the gel, FopA was mixed with native-PAGE Sample Buffer (100 mM sodium chloride, 100 mM imidazole-HCl, 4 mM 6-aminohexanoic acid, 10% glycerol, 2 mM EDTA, pH 7.0) in the volume ratio of 1:2. Electrophoresis was performed at 150 V, 10 mA using Anode Buffer (25 mM imidazole, pH 7.0) and Cathode Buffer (0.05% sodium deoxycholate, 0.01% βDDM, 50 mM Tricine, 7.5 mM imidazole, pH 7.0) in the electrophoresis chamber system (Invitrogen, Life Technologies) at 4°C.

### Determination of FopA relative molecular weight by SEC

The relative molecular weight (*M*_*r*_) of FopA was determined by size exclusion chromatography (SEC). SEC was performed as described above with minor modifications. FopA at 1.6 mg/mL in 400 μL was injected onto the SEC column and eluted at 0.5 mL/min. Molecular weight (MW) standards used were thyroglobulin (*M*_r_ 669,000), ferritin (*M*_r_ 440,000), aldolase (*M*_r_ 158,000), conalbumin (*M*_r_ 75,000) and ovalbumin (*M*_r_ 44,000) resuspended in 500 μL of 20 mM Tris, pH 7.5, 150 mM NaCl at a concentration of 3 mg/mL except for ferritin which was prepared at 0.3 mg/mL. Using the elution volume (V_e_), the elution volume parameter *K*_av_ (partition coefficient) for each MW standard was calculated and a plot of *K*_av_ versus log MW was prepared to obtain the SEC column calibration curve. FopA was eluted in its SEC purification buffer and the corresponding *K*_av_ was calculated, which was applied into the calibration curve equation to determine the *M*_r_ of FopA.

### Dynamic light scattering

Dynamic light scattering (DLS) was performed in a SpectroSizeTM302 spectrometer using a 5μL droplet of FopA at a concentration of 11 mg/mL in the SEC buffer. The mean radius distribution represented the average of 20 measurements carried out for 10 s each at 20 °C.

### Negative stain electron microscopy

For negative stain electron microscopy, FopA was freshly purified at ∼0.3 mg/mL with the purification protocol described above. Prior to sample preparation, FopA was diluted to 0.3 μg/mL (1:1000) with the SEC Buffer. FopA was applied to the glow-discharged carbon-coated 400 mesh copper grids (Electron Microscopy Sciences) and excess protein solution was removed by blotting with filter paper. The grid was immediately touched to a drop of stain solution made of 0.75% (w/v) uranyl formate and excess stain solution was removed promptly with a filter paper. The grids were let air-dry at room temperature. Data acquisition was performed using a Philips CM12 transmission electron microscope (Eyring Materials Center, Arizona State University) operated at 120 kV, and the images were obtained at 140K magnification.

### Small angle X-ray scattering experiments

Small angle X-ray scattering (SAXS) experiments were performed using an inline SEC set up at the BioCAT beamline 18ID, Advanced Photon Source (APS), Argonne National Laboratory. FopA samples were prepared in the SEC Buffer at ∼5 mg/mL, injected into the SEC column at 20 °C and eluted directly into the 1.5 mm ID quartz capillary cell at a flow rate of 0.75 mL/min. SAXS data were collected using a Pilatus3 1M (Dectris) detector at 3.5 m from the sample at an X-ray beam wavelength of 1.033 Å. The momentum transfer (*q*) measurement range was 0.004 Å^-1^ - 0.4 Å^-1^. FopA data was evaluated using the Primus program in ATSAS (40, 57). The pair distance distribution function was calculated using GNOM in ATSAS (41). Molecular weight (MW) was estimated using two methods: Porod volume (MW_Porod_) (44) and Volume of Correlation (MW_Vcorr_) (45) by GNOM in ATSAS using the DATPOROD and DATVC tools (57). MW estimation was repeated using two other alternate software: BioXTAS RAW (42) and denss.fit_data.py in DENSS software package (43). DENSS version 1.6.2 was used to reconstruct a 3D electron density map from the SAXS data. The denss.all.py script in DENSS was used to calculate twenty individual reconstructions using MEMBRANE mode and align and average the reconstructions. Fourier shell correlation was used to estimate the resolution of the map at a cutoff of 0.5 as implemented in the denss.all.py script. PyMOL (58) was used to generate images of the averaged density using the volume display mode.

### Crystallization and X-ray data collection

To identify the crystallization condition of FopA, crystallization screens were performed using the commercial crystallization kits MembFac HT™, Peg/Ion HT™, Crystal Screen HT™, Index HT™, Natrix HT™, Grid Screen Salt HT™ (Hampton Research) and MemGold MD1-39 and MemGold2 MD1-63 (Molecular Dimensions) by high throughput crystallization robot Phoenix HT (Rigaku). The initial crystallization condition of FopA (27% 2-methyl-2,4-pentanediol (MPD), 0.1 M Bis-Tris, pH 6.0, 1 mM CaCl_2_) was identified from MD1-45 MemPlus™ HT-96 screen (Molecular Dimensions). Using the above condition, FopA was crystallized at a concentration of 10 - 15 mg/mL at 18°C by the hanging drop vapor diffusion method. Crystallization drops contained equal volumes (2 µL) of reservoir solution and purified FopA. Crystallization of mature FopA was verified as follows. Approximately 10 crystals were harvested from the crystallization drop and directly transferred to a drop of reservoir solution to remove any residual proteins adhered on the crystal surface. These crystals were then transferred to a drop of SDS-PAGE Sample Buffer (supporting information), heated at 95 ° for 10 min and loaded on 12% SDS-PAGE gel. The initial crystallization condition was further optimized and yielded a slightly modified condition of 27% or 30% MPD, 0.1 M Bis-Tris, pH 6.0, 1 mM CaCl_2_ with or without the presence of 5% PEG 2000. For X-ray diffraction data collection, crystals were harvested, soaked promptly in a drop of cryo solution that was composed of the reservoir solution supplemented with 25-30% (vol/vol) glycerol and directly cryo cooled in liquid nitrogen. X-ray data collection was conducted at GM/CA beamline 23-ID-D at APS using a Pilatus3-6M detector. Data were collected at a photon energy of 12 keV at 100 K with a rotation angle of the crystals of 0.2 deg./image. Diffraction images were indexed and integrated using XDS (59)

## Acknowledgements

This research used resources of the Advanced Photon Source, a U.S. Department of Energy (DOE) Office of Science User Facility operated for the DOE Office of Science by Argonne National Laboratory under Contract No. DE-AC02-06CH11357. This project was supported by grant P30 GM138395 from the National Institute of General Medical Sciences of the National Institutes of Health. Use of the Pilatus 3 1M detector was provided by grant 1S10OD018090 from NIGMS. The content is solely the responsibility of the authors and does not necessarily reflect the official views of the National Institute of General Medical Sciences or the National Institutes of Health. We acknowledge the use of facilities within the Eyring Materials Center at Arizona State University supported in part by NNCI-ECCS-1542160.

## Supporting information captions

**S1 Fig. Amino acid sequence alignment of FopA (aa 1-393) and seven most closely related OmpA family proteins from the BLAST search**. From top to bottom, the proteins are from *Fangia hongkongensis* (FH), *Caedibacter taeniospiralis* (CT), *Fastidiosibacter lacustris* (FL), *Cysteiniphilum* sp. JM-1 (CJ), *Marinagarivorans algicola* (MA), *Alishewanella jeotgali* (AJ) and *Pseudomonas aeruginosa* (PA). The sequence numbering is for FopA. Residues that are 100% conserved are in red background with white text. For a given position, if 70% of the amino acids are identical or similar, these residues are highlighted in yellow, with the similar residues in bold.

**S2 Fig. Structure based sequence alignment of FopA-CTD (PDB ID 6U83; aa 240-393) with homologous OMPs of known structure**. The secondary structural elements of each OMP-CTD are shown above the sequences. The sequence numbering is for FopA. The OMPs used in the analysis are: *Pseudomonas* aeruginosa (PA) OprF-CTD (PDB ID 5U1H), *Escherichia coli* (EC) OmpA-CTD (PDB ID 2MQE), *Salmonella enterica* (SE) OmpA-CTD (PDB ID 4ERH) and *Klebsiella pneumoniae* (KP) OmpA-CTD (PDB ID 5NHX). Residues that are 100% conserved are in red background with white text. For a given position, if 70% of the amino acids are identical or have similar physico-chemical properties, they are highlighted in yellow, with similar residues in red characters. β-strands (black arrows), α-helices (spirals) and 310-helices (η) are indicated. Gaps are represented by black dots.

**S3 Fig. Detergent screen for solubilizing membrane embedded FopA**. Each lane in this anti-FopA immunoblot was loaded with the detergent-solubilized supernatant from an *E. coli* membrane fraction, except for the leftmost and rightmost lanes, which were loaded with MW size standards and whole cell lysate (WCL), respectively. No detergent was added to the WCL. Full-length FopA is indicated by a black arrow.

**S4 Fig. Homogeneity analysis of affinity purified FopA followed by two tandem SEC**. (A) Dynamic light scattering and (B) negative stain electron microscopy. Both analysis reveals a homogenous distribution of particles with a size range of 8-10 nm.

**S1 Table. Closest FopA homologs identified by BLAST search** ^**a**^.

^a^ Listed are all of the homologs with at least 25% sequence identity to FopA.

^b^ Percentage of query sequence residues that overlap with the reference sequence.

^c^ Expect value (E) is a parameter that describes the number of hits one can “expect” to see by chance when searching a database.

^d^ Quantitative measurement of the similarity between two sequences.

**S2 Table. FopA secondary structure composition estimated by bioinformatics and CD spectroscopy**.

**S3 Table. FopA crystals X-ray diffraction**.

**S1 File. Expression clone detail**. (A) Protein sequence. (B) Plasmid DNA sequence.

## Notes

### Competing Interest Statement

The authors have declared no competing interest.

